# Molecular mechanisms governing circulating immune cell heterogeneity across different species revealed by single cell sequencing

**DOI:** 10.1101/2021.12.17.472326

**Authors:** Zhibin Li, Chengcheng Sun, Fei Wang, Xiran Wang, Jiacheng Zhu, Lihua Luo, Xiangning Ding, Yanan Zhang, Peiwen Ding, Haoyu Wang, Mingyi Pu, Yuejiao Li, Shiyou Wang, Qiuyu Qin, Yanan Wei, Jian Sun, Xiangdong Wang, Yonglun Luo, Dongsheng Chen, Wei Qiu

## Abstract

**Background:** Immune cells play important roles in mediating immune response and host defense against invading pathogens. However, insights into the molecular mechanisms governing circulating immune cell diversity among multiple species are limited.

**Methods:** In this study, we compared the single-cell transcriptomes of 77□957 immune cells from 12 species using single-cell RNA-sequencing (scRNA-seq). Distinct molecular profiles were characterized for different immune cell types, including T cells, B cells, natural killer cells, monocytes, and dendritic cells.

**Results:** Results revealed the heterogeneity and compositions of circulating immune cells among 12 different species. Additionally, we explored the conserved and divergent cellular crosstalk and genetic regulatory networks among vertebrate immune cells. Notably, the ligand and receptor pair VIM-CD44 was highly conserved among the immune cells.

**Conclusions:** This study is the first to provide a comprehensive analysis of the cross-species single-cell atlas for peripheral blood mononuclear cells (PBMCs). This research should advance our understanding of the cellular taxonomy and fundamental functions of PBMCs, with important implications in evolutionary biology, developmental biology, and immune system disorders.

## Introduction

Peripheral blood mononuclear cells (PBMCs) are derived from myeloid and lymphoid hematopoietic systems and are mainly comprised of circulating multi-functional immune cell types, such as lymphocytes, monocytes, and dendritic cells (DCs). As the supervisor and executor of body defense, PBMCs play important roles in mediating innate and adaptive immune responses, maintaining immune homeostasis, and reflecting the real-time cellular and humoral immune state of the whole body. As such, PBMCs are widely used in the fields of immunology^1,2^, infectious diseases^3,4^, cancer^5^, vaccine development^6^, transplantation^7^ and high-throughput screening^8^. As a commonly used *ex vivo* cellular model in immunological function studies, PBMCs also play a vital role in immunological research and immunotherapy and have been used to predict biomarkers and discover potential immunotherapy targets^9,10^.

Traditional RNA sequencing (RNA-seq) provides the ability to measure average gene expression of the entire transcriptome from bulk cells, which can hide potential heterogeneity^11^. However, single-cell RNA-seq (scRNA-seq), in combination with next-generation sequencing, offers an unbiased approach to deconvolve the heterogeneity of immune cells and profile cell breadth (cell number) and depth (gene number per cell)^12^.

Due to the complexity of PBMCs, it is difficult to study the function of individual immune cells. However, advances in scRNA-seq allow comprehensive analysis of the immune system at the single-cell resolution. Thus, scRNA-seq can capture gene expression of individual immune cell types, identify new immune cell populations, reveal pathogenic immune cell subsets and transcriptional modules related to pathogenesis, and evaluate immunotherapy efficacy and response^13^. Furthermore, differential gene expression and intercellular interactions among immune cell types and samples can be evaluated^14^. In addition to humans, there have been many studies have been conducted on scRNA-seq of PBMCs in mouse models, providing data on immune cells under healthy and diseased conditions^15-17^. These studies have not only identified immune cell types, their interactions, and regulatory molecular mechanisms, but also identified potential targets for immune-related disease therapy^9,10^. scRNA-seq can be used to identify distinct immune cell types and characterize cytokine expression profiles and heterogeneity in different cell populations. In this study, we compared the cellular taxonomy of PBMCs in 12 species, revealing the conserved and divergent patterns of cellular crosstalk and genetic regulatory networks among multiple species.

## Results

### Single-cell transcriptomic profiles of PBMCs

To enable cross-species comparison of the molecular mechanisms governing immune cell heterogeneity and compositions, we first isolated the fresh PBMCs from healthy individuals of seven species: i.e., cat (*Felis catus*), dog (*Canis lupus familiaris*), rabbit (*Oryctolagus cuniculus domesticus*), hamster (*Mesocricetus auratus*), deer (*Cervus nippon*), goat (*Capra aegagrus hircus*), and pigeon (*Columba livia domestica*). Using high-throughput single-cell RNA-seq and filtering of doublets and low-quality cells (see Methods), we obtained high-quality scRNA-seq data of 50□478 cells. We next integrated five publicly available PBMC scRNA-seq datasets: i.e., human^18^, tiger^19^ monkey^20^, mouse^21^, and zebrafish^22^. After filtering doublets and low-quality cells, single-cell transcriptome data of 27□479 cells were obtained for further analyses (**Fig. 1A**).

**Figure 1.**
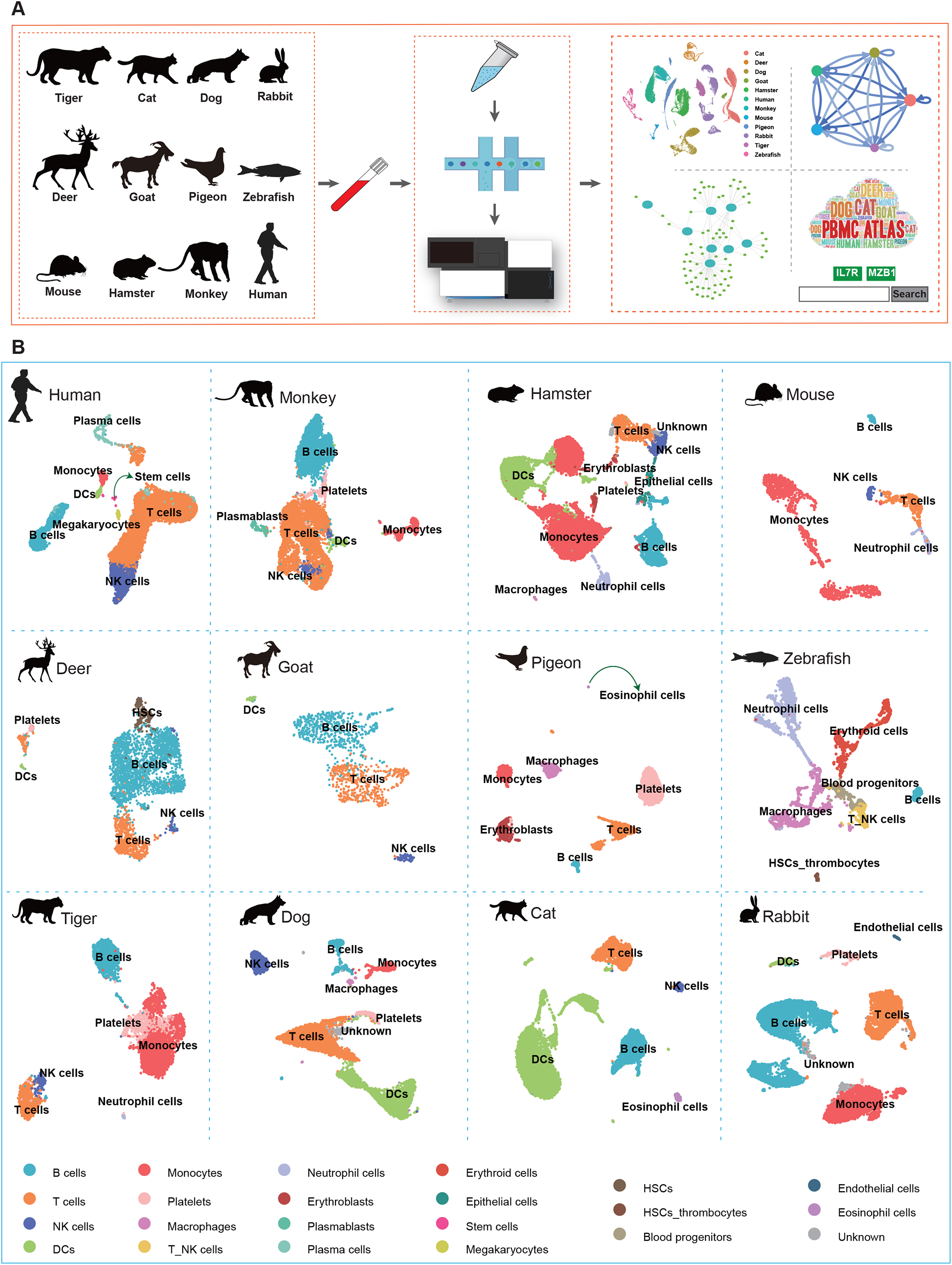

We first humanized the homologous genes for all nonhuman species and performed unsupervised clustering using the top variable genes. We identified five main types of immune cells, i.e., T cells, B cells, natural killer (NK) cells, monocytes, and DCs based on the specific expression of cell-type marker genes (**Fig. 1B**). Cell type identity was also confirmed using Gene Ontology (GO) functional enrichment analysis of differentially expressed genes (DEGs) corresponding to clusters. For example, in the cat, B cells were annotated based on the high expression of markers *CD19, IRF8*, and *MS4A1*; T cells were characterized by the specific expression of *TCF7* and *TAPBPL*; NK cells were identified by enrichment of *CCL5, GZMA*, and *KLRF1*; and DCs were characterized by enrichment of *TREM1, TKT*, and *SOD2* (**Fig. 2A**,**B**). In the dog, B cells were annotated based on the high expression of markers *CD19, BLNK, FCRLA*, and *MS4A1*; T cells were characterized by the specific expression of *CD4, SOD2, ITM2B*, and *SELL*; NK cells were identified via enrichment of *GZMB*; DCs were confirmed by enrichment of *MAFB* and *RETN*; and monocytes were identified by the expression of *IL-7R, TCF7*, and *CCR7* (**Fig. 2C**,**D**). The specific expression patterns of these molecules successfully identified the different cell types and provided a molecular basis for exploring the physiological functions of the respective immune systems. In both cats and dogs, functional analysis of the five main immune cells in PBMCs indicated that they were primarily related to regulation of innate immune response, regulation of immune effector process, neutrophil activation involved in immune response, immune response-activating signal transduction, and immune response-activating cell surface receptor signaling pathway.

**Figure 2.**
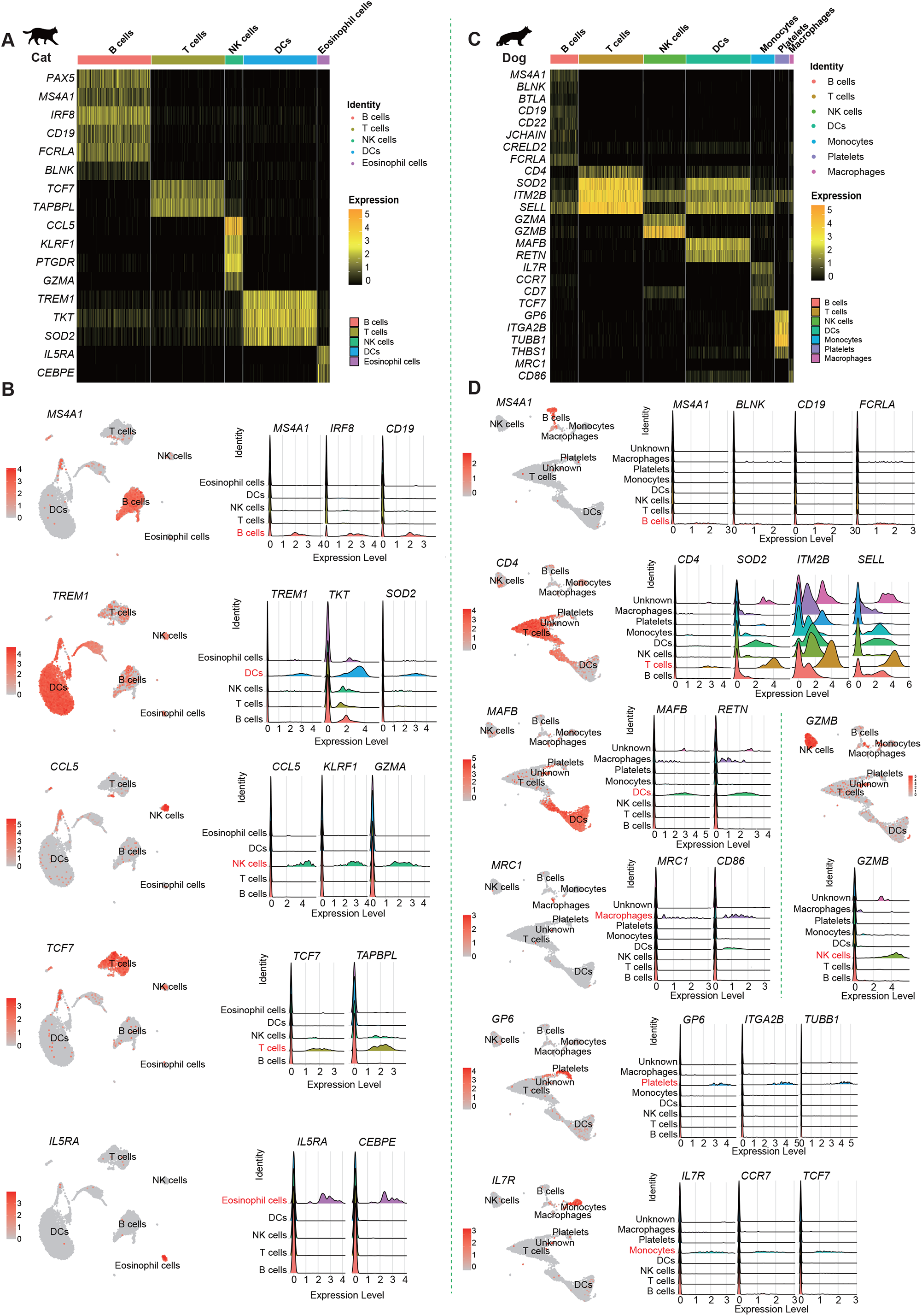

### Conservation of PBMC connectomes

To identify potential cellular interactions, we constructed a ligand receptor-mediated communication network of the above five immune cell types for each species using the Connectome R package^23^. In general, source connectome topology of the different species was very similar (**Fig. 3A**). Based on the interaction network data, we further analyzed the relationship between ligand-receptor pairs in the different cell types. For example, CD4OLG was identified as a ligand of T cells, and was able to interact with its receptors CD40, TRAF3, and ITGB2 on B cells (**Fig. 3B**). For target analysis of network cross-plot centrality, these communication pairs were generally classified into seven signaling modalities (i.e., tumor necrosis factor (TNF), NOTCH, matrix glycoproteins, intracellular trafficking, interleukins, complement, and uncategorized) (**Fig. 3C**). In addition, T cells interacted frequently with other cell types, while the NK cells were relatively less active (**Fig. 3C**,**D**).

**Figure 3.**
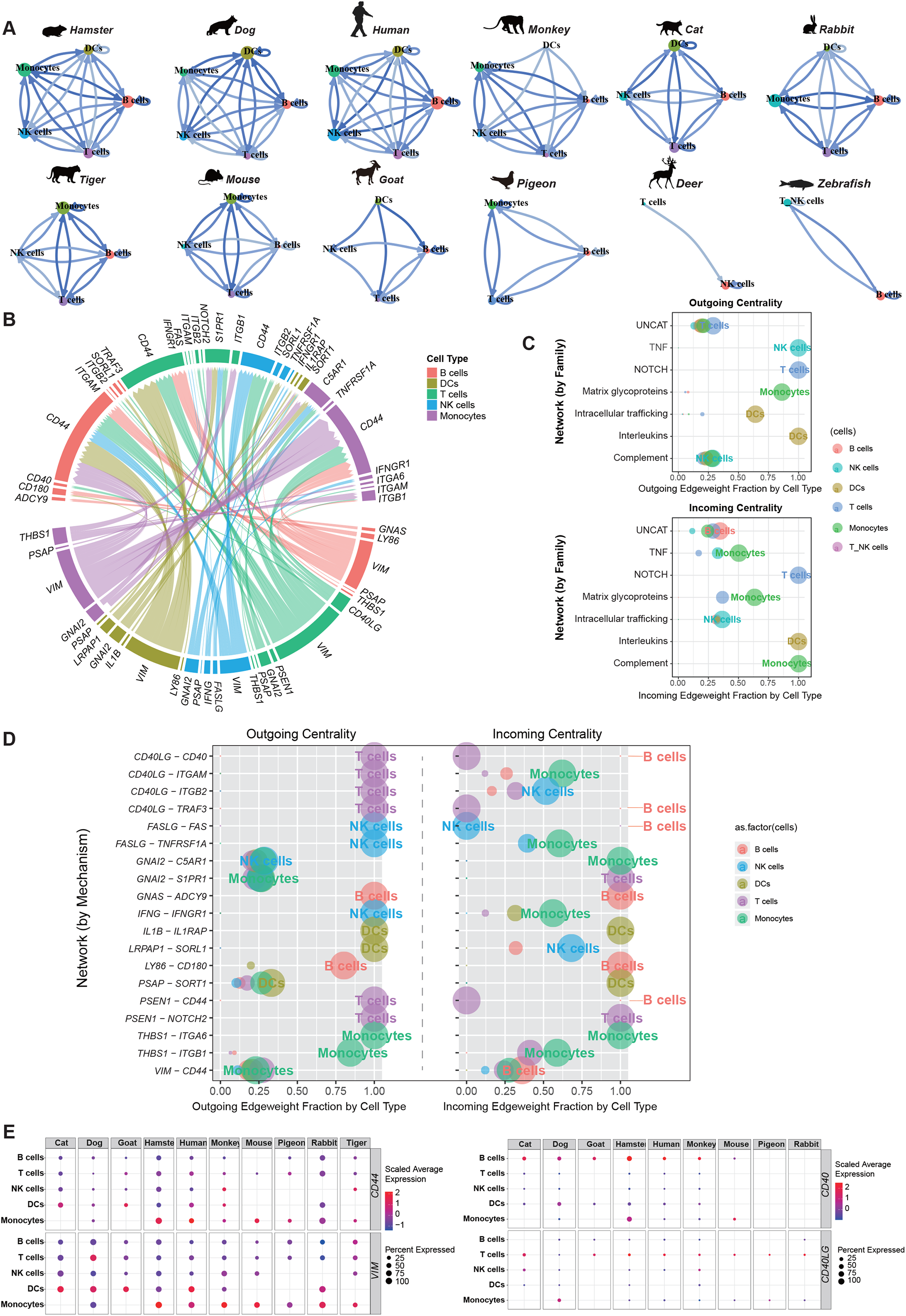

We further identified connectivity among immune cells, which may correspond to common ancient communication signals. In total, 214 pairs of cell-cell connections were conserved among the five main immune cells in the 12 species. Among them, there were 19 ligand-receptor pairs showing pan-conserved immune cell interactions. In addition, ligands related to the *VIM* gene appeared frequently, and receptors related to the *CD44* gene appeared most frequently, with most belonging to the uncategorized signaling modality (**Fig. 3D**). In addition to immune cell crosstalk, we explored the conservation of cell-cell crosstalk in different species. Several interactions between immune cells in PBMCs, such as the VIM ligand and CD44 receptor, were commonly expressed in the DCs and T cells of humans, monkeys, hamsters, dogs, cats, rabbits, and goats. As for the CD40LG and CD40, CD40LG and TRAF3, CD40LG and ITGB were commonly expressed in T cells-B cells among humans, monkeys, hamsters, mice, tigers, dogs, and cats. The above four pairs belonged to the uncategorized signaling modality. The GNAI2-C5AR1 ligand-receptor pair was expressed in the monocytes of six species (human, monkey, hamster, mouse, rabbit, and dog), and categorized as complement signaling modality. Additionally, the PSAP-SORT1 ligand-receptor pair was expressed in the DCs of five species (human, hamster, cat, rabbit, and dog) and belonged to the intracellular trafficking signaling modality (**Fig. 3E**).

### Conservation of PBMC regulomes

To explore the regulatory mechanisms underlying immune system development in the light of evolution, the PBMC genetic regulatory networks were predicted for the 12 species. The most conserved genes that encoded transcription factors (TFs) were *ARID5B, MEF2C, PAX5, NCOR1, FOS, CSDE1, SUB1*, and *CCDC88A* (**Fig. 4A**). Subsequently, we analyzed the regulatory network of the five immune cells. A variety of TF-target interactions conserved in at least four species were identified (293 in T cells, 324 in B cells, 108 in DCs, 94 in NK cells, and 194 in monocytes) (**Fig. 4B**). Enrichment of GO terms for predicted target genes indicated that the regulatory functions of these TFs were closely related to immune response processes (**Fig. 4C**). Interestingly, several regulators for B cells, T cells, and DCs were active in the genetic regulatory network of the corresponding cell types, in line with their expected regulatory functions (**Fig. 4C**). Many regulatory circuits were highly conserved in each cell type among the multiple species (**Fig. 4C**). Specifically, in B cells, *MEF2C* (regulatory gene of *LYN, ABRACL, ARID5B, CANX, CCT5, HSP90B1, PAX5, PLEK, PNISR, RALGPS2, SNW1, VIM*, and *PAN3*) was conserved in at least 10 species. *PAX5* and its target genes (*LYN* and *VCP*) were conserved in at least six species, and were mainly related to the differentiation, activation, proliferation, and receptor signaling pathway of B cells. In T cells, the conserved regulatory relationships between *NCOR1* and its target genes (*PTPRC, ARHGAP15, IQGAP1, RPL27, SCAF11*, and *YTHDC1*), *TCF7* and its target gene (*LEF1*), and *FOS* and its regulatory genes (*PTPN6, TMBIM6, RPL27, HERPUD1*, and *VIM*) were conserved in at least six species, and were mainly involved in the differentiation, activation, proliferation, and receptor signaling pathway of T cells. In DCs, *CCDC88A* and its target genes (*PSMA3, GOLGA4, LGALS1, PDIA3*, and *SWAP70*) and *PLEK* and its target genes (*ANXA1* and *SGK1*) were conserved in at least five species and were mainly related to antigen processing and presentation. In NK cells, *SUB1* and its target gene (*NDUFA8*) were conserved in at least six species. In monocytes, *PLEK* and its target genes (*B2M, CTLA*, and *LGALS3*) and *ZEB2* and its target gene (*NPC2*) were conserved in at least six species.

**Figure 4.**
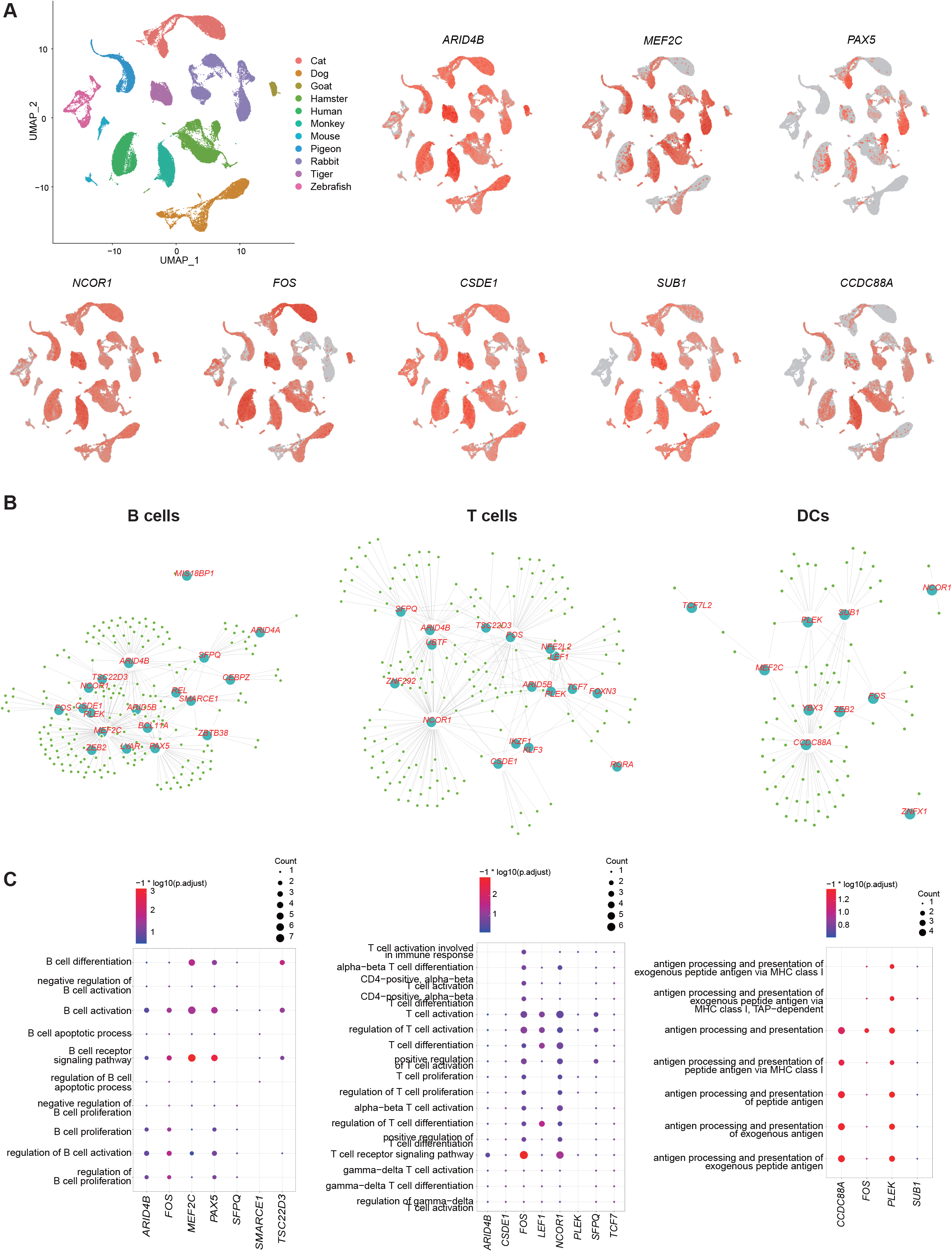

## Discussion

As the most often used cell model in immunological studies, PBMCs can reflect the dynamic changes in circulating innate and adaptive immune systems and play a vital role in immunotherapy. Here, we used scRNA-seq and atlas analysis to elucidate the heterogeneity of PBMCs, and investigated the conserved cell-cell communications and genetic regulatory networks of major immune cell types across 12 different species.

### Cellular heterogeneity and compositions of PBMCs

We surveyed the PBMC atlas by scRNA-seq for comprehensive analysis. In total, 77□957 cells were derived from the 12 species. After cell clustering analysis, we identified five major immune cell types for each species, including NK cells, B cells, DCs, monocytes, and T cells. The divergences of immune cells across species were also investigated in previous study^24^.

### Conserved cellular crosstalk between immune cells

#### 1) VIM-CD44 in DCs-T cells

We analyzed cell-cell interactions from the classical immune response and focused on DCs-T cells and T cells-B cells, which are involved in antigen presentation processing and adaptive immune response. The top interactions between *VIM* and *CD44* in DCs-T cells were identified among 214 pairs of conserved cell interactions in the 12 species. VIM-CD44 is a highly conserved cell-cell crosstalk pair, especially among ligand-receptor pairs in DCs-T cells. *VIM* is expressed in lymphocytes and can interact with other proteins for intercellular signal transduction and can be released as an antigen component of pathogen infection^25,26^, with bacterial and viral pathogens able to attach to this protein on the host cell surface^27^. *CD44* is up-regulated in activated lymphocytes and is involved in various cellular functions, including activation, recirculation, and homing of T-lymphocytes (when stimulated by CD44, T cells are activated and IL-2 production is elevated under CD44 stimulation), as well as hematopoiesis, inflammation, and response to bacterial infection^28^. In our study, the VIM-CD44 interactions were highly conserved among multiple species. VIM in DCs cooperates with CD44 in T cells, which promotes the antigen presentation and activation of autoreactive T cells. Additionally, the binding of VIM and CD44 activates T cells, triggers activation of a series of possible effector genes, and activates signaling pathways transduction^29^. However, how this signal is transduced and participates in immune responses and host defense requires further study in immune diseases.

#### 2) CD40LG-CD40 in T cells -B cells

The commonly conserved ligand-receptor pair in T cells-B cells was identified as CD40LG-CD40 in our study. CD40LG is predominantly expressed in activated CD4^+^ T cells and binds to its ligand (CD40) on the surface of B cells, thereby influencing B cell function^30,31^. This pair also co-stimulates T-cell proliferation and cytokine production^32^ and enhances the expression of IL-4 and IL-10^32^. *CD40LG* deficiency is a severe primary immunodeficiency caused by mutations in the *CD40L* gene, which can lead to T cell impairment, B cell defects, and susceptibility to opportunistic pathogens^33^. Mutation of the *CD40* gene can result in type 3 hyper-IgM immunodeficiency, characterized by an inability to undergo isotype switching, an inability to mount an antibody-specific immune response, and a lack of germinal center formation^34^. Clinical trials have evaluated novel therapeutic approaches targeting the CD40-CD40LG pathway based on T cell-B cell interactions for autoimmune diseases^35^.

### Distinct conserved regulatory networks in different immune cells

#### 1) CCDC88A and its target genes in DCs

Lastly, we investigated the conserved regulatory networks in different immune cells among the 12 species. The coiled-coil domain containing 88A (*CCDC88A*) gene was conserved within five species. Its encoded protein is a kind of coiled-coil domain containing Girdin family proteins that are activated by Akt and necessary for cytoskeleton remodeling and cell migration^36-38^. CCDC88A accumulates in cell protrusions and contributes to the formation of membrane protrusions and cell migration and invasion^39,40^, in line with the structural characteristics and functional demands of DCs. In addition, *PSMA3* a target gene of *CCDC88A*, which encodes proteasome, is essential for the generation of a subset of major histocompatibility complex (MHC) class I-presented antigenic peptides, as well as for the maturation of DCs^41^. The GIV protein of *CCDC88A* is most highly expressed in DCs and macrophages and participates in the inhibition of proinflammatory signaling via Toll-like receptors (TLRs)^42^. These developmental and functional characteristics of DCs may provide new pathways to restore immune tolerance and inhibit self-antigen presentation processing.

#### 2) NCOR1, TCF7, and their target genes in T cells

Nuclear receptor corepressor 1 (*NCOR1*) was conserved in the T cells of 11 species and functioned as a transcriptional corepressor by connecting chromatin-modifying enzymes with gene-specific TFs. *NCOR1* plays an important role in controlling positive and negative selection of thymocytes during T cell development^43^. Furthermore, *NCOR1* is considered a novel regulator of immune tolerance and immune cell development^44^, and shapes the transcriptional landscape, influences the direction of CD4^+^ T cell differentiation, and controls Th1/Th17 effector functions^45^. In addition, its target gene *PTPRC* (protein tyrosine phosphatase receptor type C), also known as CD45, is essential for T cell antigen receptor-mediated activation, and its downstream regulatory imbalance can result in autoimmunity^46,47^. As another crucial regulatory gene, *TCF7* is predominantly expressed in T cells and plays a critical role in T cell development^48,49^. Its encoded protein, T cell factor-1 (TCF-1), belongs to the T-cell factor/lymphoid enhancer-binding factor family. This encoded protein is highly expressed in naive CD8^+^ T cells but is down-regulated after differentiation into effector CD8^+^ T cells, and is necessary for the formation of central CD8^+^ T cell memory in response to infection^50-52^. Silencing of *Tcf1* facilitates effector CD8^+^ T cell differentiation^51^, and knockout of *TCF7* in mice results in impaired T-lymphocyte differentiation^53^. In addition, its target gene *LEF1*, which encodes lymphoid enhancer binding factor 1, can bind to functionally important sites in the T cell receptor-α enhancer. This is critical for the maturation and development of IL17A-producing T cells, with its imbalance downstream potentially resulting in autoimmunity^47,54^.

#### 3) MEF2C, PAX5, and their target genes in B cells

Our results showed that myocyte enhancer factor 2C (*MEF2C*) was conserved in the B cells among 11 species. *MEF2C* binds the active regulatory region to the V(D)J gene in mouse B cell progenitors and human B lymphoblasts, which is essential for lymphatic fate determination^55,56^. MEF2C has a highly conserved MADS box and MEF2 domain, which contribute to B cell homeostasis^57,58^. In addition, MEF2C and early B cell factor-1 together form a co-regulator, which targets and regulates a subset of B cell-specific genes ^55^. Various animal models also show that *MEF2C* is important in myeloid leukemia. Mutations in *MEF2C* are often found in patients with B cell lymphoma, and these mutations are involved in the pathogenesis of abnormal B cell proliferation^57,59,60^. *LYN* is the target gene of *MEF2C*, participating in the regulation of B cell differentiation, proliferation, survival, and apoptosis, and plays an important role in maintaining immune self-tolerance^61^. It also acts downstream of B cell receptors via the down-regulation of signaling pathways. As another crucial regulatory gene of B cells, paired box protein 5 (*PAX5*) is necessary for the differentiation of lymphoid progenitor cells into B lymphocyte lineage^62^. *PAX5* regulates transcriptional reprogramming processes by restricting uncommitted progenitors to the B cell pathway, promoting V(H)-DJ(H) recombination, inducing B cell receptor signaling, and facilitating development to the mature B cell stage^62,63^. However, *PAX5* inhibition is not necessary for stable plasma cell development and antibody secretion, even though it is essential for immunoglobulin G (IgG) production and long-lived plasma cell increase^64^. Thus, the role of *PAX5* in plasma cell differentiation needs to be further investigated. *LYN* is also a target gene of *PAX5*, and studies show that both are related to B cell development^65,66^, however, the complex and sophisticated function for evolutionary evidence in regulating B cell development needs to be further investigated. Collectively, we identified highly conserved regulons in the PBMCs communities of different animal species. Identifying conserved key genes and exploring their functions in multiple species will help improve our understanding of the development, maturation, proliferation, activation, differentiation of immune cells.

In this study, we produced the first comprehensive PBMC atlas of 12 species, which holds significance for immunological research. We systematically studied the gene expression profiles and molecular characteristics of each cell type and compared them across species at single-cell resolution. We also identified key genes and highly conserved cell-cell interactions that play important roles in regulating development and immune response. The PBMCatlas website was constructed, which provides an accessible approach to explore the different species datasets. These results provide a systematic resource for understanding the immune cell diversity and as well as insights into the molecular mechanisms governing conservation of PBMCs across species.

### Limiations of the study

A few limitations of this study were also addressed, which should be investigated by further study. First, the conservation of homologous genes in all non-human species were calculated to identify the distinct cell types of PBMCs in 12 different species. However, the non-traditional species (deer, rabbit etc.) showed the low ratios of homologous genes of these species to human, resulting in the lower diversity in cell types of PBMCs for these species. Second, when comparing of cell-type specific gene expression across species, the effects of age, sex, and physiological conditions were not considered in this study. Nevertheless, major immune cell types were identified in all these species, and the cellular heterogeneity and compositions of PBMCs were characterized. The connectome analysis might be influenced by potential species differences in ligand-receptor interaction relationships or homolog conversion. Last, the aim of this study was to focus on the single-cell altas of PBMCs from multiple species, even though mRNA and protein expression in immunce cells show discrepant by different methods^67-69^. In addition, the datasets of PBMCs for 12 species were obtained from different platforms, and differences caused by sample processing, scRNA-seq technical bias, and batch effects could impact the cells capture and cell type classifications, resulting in biological differences. Most importantly, despite these limitations, this study provides a key resource of PBMCs for understanding the immune cell type diversity ae well as insights into the developmental and evolutionary biology of the circulating immune system across species.

## Material and methods

### Blood samples collection and ethics statement

Blood samples were obtained from seven animals, including: goat (*Capra aegagrus hircus*), cat (*Felis catus*), hamster (*Mesocricetus auratus*), dog (*Canis lupus familiaris*), pigeon (*Columba livia domestica*), rabbit (*Oryctolagus cuniculus domesticus*), and deer (*Cervus nippon*). The deer blood samples with related genome assembly were provided by the Institute of Special Animal and Plant Sciences (ISAPS) of the Chinese Academy of Agricultural Sciences. Other blood samples were obtained from farmers’ markets with permission from the BGI Ethics Committee. The gene expression matrices for five other species (human^18^, tiger^19^, monkey^20^, mouse^21^, zebrafish^22^) were obtained published datasets. The Institutional Review Board on Ethics Committee of BGI reviewed and approved this study (NO. BGI-IRB A20008, BGI-IRB A20008 T1).

### Peripheral blood mononuclear cells (PBMCs) processing

All operations were performed under sterile conditions. Peripheral blood samples (3 mL) were collected into an EDTA anticoagulant tube, gently reversed 4-6 times, and fully mixed, and placed at room temperature. Whole blood was diluted with 3 mL of phosphate-buffered saline (PBS) and transfered in to a 15 mL centrifuge tube. After this, 6 mL of Histopaque-1077 (Cat. No. 10771-6×100ml) was slowly added into the 15 mL centrifuge tube, followed by density gradient centrifugation methods to collect peripheral blood mononuclear cells (PBMCs).

### Single cell RNA-seq library construction and sequencing

The goat, deer and pigeon PBMCs underwent library construction using DNBelab C Series Single-Cell Library Prep Set (MGI) and sequenced with BGISEQ-500 in China National GeneBank (CNGB). As for the cat, dog, rabbit and hamster PBMCs underwent library construction using a 10X Chromium Next GEM Single cell 3’ Reagent Kits v3.1 following the guidelines provided by the manufacturer and sequenced using NOVAseq 6000 sequencing platform of Illumina.

### Cross-species homologous gene conversion and single-cell RNA-seq data processing

We refered to the previous methodological section for cross-species homologous gene transfer and single-cell RNA-seq data processing^70^.

### Cell type annotation

For annotation of the self-produced datasets, we used several classic cell type markers from the CellMarker database^71^. The four published datasets used cell type makers from the published article mentioned. The annotation results are presented in Figure 1b using UMAP feature plots.

### Differentially expressed genes (DEGs) and Gene Ontology (GO) term enrichment analysis

All differentially expressed genes (DEGs) for each cell type were identified using the FindAllMarkers function in Seurat with only.pos set to T, min.pct = 0.1, logfc.threshold = 0.25. The hypergeometric test implemented in the clusterProfiler^72^ package with the compareCluster function (ENTREZID∼celltype, fun=“enrichGO”, “org.Hs.eg.db”, pvalueCutoff = 0.05) was used to carry out Gene Ontology (GO) term enrichment analysis.

### Conserved cellular communication analysis

We applied the Connectome (https://github.com/msraredon/Connectome) R package^23^ for analysis. All ligands and receptors were downloaded from the FANTOM5 database^73,74^. Firstly, the five major immune cells (B cells, T cells, NK cells, DCs, Monocytes) were extracted from the annotated datasets of the 12 species for further analysis. Then connectome networks were then constructed according to the expression of ligands and receptors.

### Conserved TF-target interaction analysis

We applied the the GENIE3^75^ R package for analysis using data from 11 species (cat, dog, goat, hamster, human, monkey, mouse, pigeon, rabbit, tiger, and zebrafish). The human TF list was downloaded from animalTFDB3.0^76^. The igraph R package^77^ was used to visualize representative regulatory TFs networks.

## Acknowledgements

We thank the China National GeneBank for producing the sequencing data. This work was supported by the National Natural Science Foundation of China (#81771300, #81971140), the Norman Bethune Foundation (#2020009) and Natural Science Foundation of Guangdong Province (#2020A1515010053).

## Author Contributions

Wei Qiu, Dongsheng Chen, Yonglun Luo, Xiangdong Wang and Jian Sun conceived and designed the study. Xiran Wang, Jiacheng Zhu and Lihua Luo prepared the samples. Chengcheng Sun, Yanan Zhang and Mingyi Pu performed all scRNA-seq experiments. Chengcheng Sun, Zhibin Li, Xiangning Ding, Peiwen Ding, Haoyu Wang, Yuejiao Li, Shiyou Wang, Qiuyu Qin, and Yanan Wei analyzed and interpreted the data. Zhibin Li and Fei Wang wrote the manuscript. Yonglun Luo, Wei Qiu, Dongsheng Chen, and Xiangdong Wang revised the manuscript. All authors read and approved the final version of the manuscript.

## Data Availability Statement

The data that support the findings of this study were deposited in the CNSA (https://db.cngb.org/cnsa; accession number: CNP0002177).

